# Acute and chronic alcohol modulation of extended amygdala calcium dynamics

**DOI:** 10.1101/2023.10.10.561741

**Authors:** Alison V. Roland, Tzu-Hao Harry Chao, Olivia J. Hon, Samantha N. Machinski, Tori R. Sides, Sophia I. Lee, Yen-Yu Ian Shih, Thomas L. Kash

## Abstract

The central amygdala (CeA) and bed nucleus of the stria terminalis (BNST) are reciprocally connected nodes of the extended amygdala thought to play an important role in alcohol consumption. Studies of immediate-early genes indicate that BNST and CeA are acutely activated following alcohol drinking and may signal alcohol reward in nondependent drinkers, while increased stress signaling in the extended amygdala following chronic alcohol exposure drives increased drinking via negative reinforcement. However, the temporal dynamics of neuronal activation in these regions during drinking behavior are poorly understood. In this study, we used fiber photometry and the genetically encoded calcium sensor GCaMP6s to assess acute changes in neuronal activity during alcohol consumption in BNST and CeA before and after a chronic drinking paradigm. Activity was examined in the pan-neuronal population and separately in dynorphinergic neurons. BNST and CeA showed increased pan-neuronal activity during acute consumption of alcohol and other fluid tastants of positive and negative valence, as well as highly palatable chow. Responses were greatest during initial consummatory bouts and decreased in amplitude with repeated consumption of the same tastant, suggesting modulation by stimulus novelty. Dynorphin neurons showed similar consumption-associated calcium increases in both regions. Following three weeks of continuous alcohol access (CA), calcium increases in dynorphin neurons during drinking were maintained, but pan-neuronal activity and BNST-CeA coherence were altered in a sex-specific manner. These results indicate that BNST and CeA, and dynorphin neurons specifically, are engaged during drinking behavior, and activity dynamics are influenced by stimulus novelty and chronic alcohol.

## Introduction

The bed nucleus of the stria terminalis (BNST) and central amygdala (CeA) are two nodes of the extended amygdala that orchestrate emotional and behavioral responses to a variety of physiological stimuli. These include aversive stimuli, such as pain and predator threat, and appetitive stimuli, such as food, drugs of abuse, and social interactions (1,2). The BNST and CeA are anatomically parallel structures, with extensive overlap in inputs as well as projection sites and similar cellular composition (3,4). Except for a sparse population of glutamatergic cells, both regions are predominantly GABAergic and express a diverse array of neuropeptides, including somatostatin, corticotropin-releasing factor (CRF), and dynorphin, among others (5). Human and non-human primate neuroimaging data indicate that BNST and CeA are functionally connected (6,7), and reciprocal anatomical connections have been demonstrated in primates and rodents (8–12). Consistent with roles in regulating both appetitive and aversive behaviors, BNST and CeA exhibit parallel output pathways with opposing functional roles. For example, in the dorsal BNST, laterally located CRF neurons projecting to the lateral hypothalamus (LH) are activated by aversive stimuli and promote avoidance, while medially located LH-projecting cholecystokinin neurons are activated by rewarding stimuli and promote approach behavior (13). In a similar vein, the BNST exhibits complementary ventral tegmental area (VTA) outputs distinguished by expression of either GABA or glutamate, which convey reward and aversion, respectively (14).

The extended amygdala has been extensively studied with respect to alcohol drinking, where it is thought to play a role in both the intoxication and withdrawal stages of the addiction process (15). Initial, nondependent alcohol consumption is associated with increased c-fos expression in BNST and CeA (16–19), and alcohol acutely modulates neurotransmission in these regions (20). BNST or CeA lesion or local pharmacology can alter alcohol intake in nondependent animals (21–25). These regions are posited to contribute to the reinforcing effects of alcohol consumption through projections to classical reward systems including the LH and VTA (15). Accordingly, silencing of BNST inputs to the VTA reduces binge drinking (26,27). Following chronic high-level alcohol intake, plasticity in extended amygdala leads to reduced reward and increased stress system engagement, promoting drinking through negative reinforcement mechanisms (28). Consistent with this idea, CeA inactivation reduces drinking in dependent animals (21), and inhibition of CeA projections to BNST reduces alcohol drinking and somatic symptoms of withdrawal (29).

Plasticity in neuropeptide systems contributes to functional changes across the addiction cycle, and the dynorphin/KOR system in the extended amygdala is strongly implicated in alcohol consumption. Dynorphin is thought to drive a dysphoric state during alcohol withdrawal (30) but may also be activated during earlier stages of drinking (31). In the CeA, acute enhancement of KOR signaling increases binge drinking; conversely, KOR antagonism, KOR or dynorphin genetic deletion, or chemogenetic inactivation of dynorphin neurons in the CeA reduces binge drinking (32). In the BNST, direct infusion of a KOR antagonist reduces binge alcohol consumption and blocks the effects of a systemic KOR agonist to increase drinking (33). KOR antagonism in either BNST (34,35) or CeA (36) also reduces drinking following the development of alcohol dependence.

Despite evidence for acute engagement of the BNST and CeA, and dynorphin neurons specifically, during alcohol consumption, the dynamics of these responses in vivo are unknown. Here we assessed these dynamics in real time using fiber photometry and the genetically encoded calcium indictor GCaMP. GCaMP6s was virally expressed and optical fibers implanted in contralateral BNST and CeA to enable simultaneous assessment of pan-neuronal responses of these regions in individual mice during acute alcohol drinking. To assess how chronic alcohol exposure alters calcium responses, recordings were made before and after exposure to a three-week continuous access alcohol paradigm. We also utilized a cre-dependent GCaMP6s in preprodynorphin-cre (pdyn-cre) mice to examine the activity of dynorphin neurons in these regions under the same conditions. BNST and CeA are known to exhibit coordinated increases in activity during exposure to anxiety-provoking stimuli; however, it is unknown if these regions coactivate during consummatory behavior or if alcohol changes their coactivation, although evidence has suggested altered connectivity between these regions following alcohol exposure (29,37). Thus, we also assessed BNST-CeA signal coherence during drinking behavior and following chronic alcohol exposure.

## Methods

### Mice

Male and female preprodynorphin-cre and mice were bred in-house at the UNC School of Medicine, and wild-type C57BL/6J mice were obtained from Jackson Laboratories. All mice were maintained on 12:12 h reverse light cycle (lights off at 0700) with food (Isopro RMH 3000, LabDiet, St. Louis, MO) and water available ad libitum outside of specified experimental water deprivation periods. All experiments were approved by the UNC School of Medicine Institutional Animal Care and Use Committee (IACUC) in accordance with the NIH guidelines for the care and use of laboratory animals.

### Stereotaxic surgery

Adult mice (7-8 weeks of age) were microinjected with 250 nL AAV5-hsyn-GCaMP6s (UPenn Vector Core, Philadelphia, PA) or AAV5-hsyn-flex-GCaMP6s into the BNST (AP: +0.3, ML: +/-0.9, DV: -4.35) and contralateral CeA (bregma: AP: -1.2, ML: +/-2.9, DV: -4.7) using a Hamilton Neuros syringe (Hamilton Company, Reno, NV), with mice counterbalanced for side placement. Virus was delivered at a rate of 100 nL/min and the syringe left in place for 5 minutes following injection to allow for viral diffusion. Fiber optic cannulae (200 μm, 0.37 nA, Neurophotometrics, San Diego, CA) were then implanted in the BNST and CeA using the same coordinates as for viral injections and fixed in place using C&B-Metabond (Parkell, Inc., Edgewood, NY). Mice were transferred to single housing one week after surgery and allowed a total of four weeks of recovery in the home cage prior to experimentation.

### Fiber photometry

Fiber photometry recordings were made using previously described methods (38). Briefly, using a commercially available system from Neurophotometrics, Inc., GCaMP6s signals were recorded from BNST and CeA simultaneously using a multi-branch patch cord. Alternating pulses of 415 nm and 470 nm light (40 Hz, 50 μW) were bandpass filtered and focused onto a multi-branch patch cord by a 20x objective. Simultaneous video recordings were obtained using a camera mounted above the bottle entry port.

One week prior to recordings, mice were habituated to patch cords for 15 minutes and brief recordings obtained to confirm signal quality. Because tethering is mildly aversive and tends to reduce consummatory behavior, mice were water deprived overnight for 12-18 hours prior to drinking experiments to encourage fluid consumption (water removed at 2100, with recordings made between 0900 and 1500 the following day). A subset of recordings was made without water deprivation for comparison. Test sessions were 10-20 minutes long. Following a baseline period of two minutes, a bottle containing the test fluid was placed on the cage and the animal allowed to freely consume the tastant. After several drinking bouts were observed, or after 10 minutes if no drinking occurred, the bottle was removed. In some cases, multiple fluids (up to 3) were tested sequentially in the same recording session. Test fluids consisted of ethanol (15% v/v), sucrose (5%, Fisher BP220-212), quinine (100 μM, Sigma Q1125), or saccharin (0.1%, Sigma 47839) dissolved in tap water, or tap water alone. In palatable chow test sessions, a single pellet of high-fat chow (60% fat, Research Diets D12492, New Brunswick, NJ) was added to the home cage.

### Continuous Access Drinking

Baseline fiber photometry recordings during drinking tests were completed over a period of three weeks, after which mice were provided three weeks of continuous access (CA) to 20% w/v alcohol and water. Although higher than the concentration tested in photometry experiments, this concentration was used to align with prior studies in our laboratory employing the CA drinking paradigm, which typically generates stable but not escalating levels of intake (39). Mice were weighed and bottles refilled weekly, and bottles were weighed daily Monday through Friday. An empty cage was mounted with a water and alcohol bottle to determine fluid loss due to drip and bottle positioning. Daily water and alcohol intake for each mouse (reported in g/kg) was determined. At the end of the alcohol access period, fiber photometry recordings were made in the home cage approximately 24-30 hours after final removal of the alcohol bottle. Following a 2-minute baseline recording, calcium activity in BNST and CeA was recorded during consumption of the following tastants, in order: 15% alcohol, 5% sucrose, water, and high-fat chow. If an animal did not consume the tastant after 5 minutes, it was removed and the next tastant offered. A final recording session was conducted 3-4 weeks (range: 22-28 days) post completion of continuous access to assess calcium activity during alcohol consumption at a later timepoint post continuous access.

Following completion of photometry recordings, mice were anesthetized with a terminal injection of avertin and transcardially perfused with 4% PFA in PBS. Brains were removed, postfixed overnight in 4% PFA, and then transferred to PBS until sectioning. 45 μM sections containing BNST and CeA were mounted on slides, coverslipped with Prolong Gold Antifade Reagent with DAPI (Cell Signaling Technology, Danvers, MA), and imaged on a Keyence BZ-X800 microscope (Keyence, Itasca, IL) for verification of virus and fiber placements.

### Data Analysis

Photometry data were analyzed using a custom MATLAB script. Signals collected at 415 and 470 nm were deinterleaved, and the background fluorescence was subtracted from each trace. Traces were lowpass filtered at 2 Hz and fit to a biexponential curve that was subtracted from each trace to correct for baseline drift. dF/F (%) was calculated as (raw signal-fitted signal)/(fitted signal) and Z-scored. The 415 nm signal was fit to the 470 nm signal using non-negative robust linear regression and then subtracted to yield the motion-corrected 470 signal. Drinking and eating bouts were manually scored, and bouts ≥ 5s in duration were used for analysis. Consecutive bouts were included only when the interval between bouts was ≥ 10s.

Statistical analyses were performed using GraphPad Prism 9 Software (La Jolla, CA). For a given tastant/stimulus, bouts from individual mice were averaged, and statistics were computed based on the group n of 13-16 mice per genotype (5-11 per sex). To assess changes in activity time-locked to consumption bouts, the mean Z score (relative to the entire recording trace) for the 5 s before bout onset was compared to the 5 s following bout onset (2s for quinine, for which drinking bouts were shorter). Two-way ANOVA with Tukey’s multiple comparison testing was used to assess sex differences; when no sex differences were detected, data from both sexes were combined and compared using paired t-testing. To compare GCaMP activity during alcohol consumption pre-CA and at two timepoints post-CA (24 hours and 3 weeks), we used mixed-model ANOVA with Tukey’s multiple comparison testing due to missing values because not all subjects drank alcohol during both post-CA recording sessions.

To determine the BNST-CeA spectral coherence, we first used Morlet wavelet transform to decompose the signal into the analytic signal according to previous methods (40). We then used the analytic signal to calculate spectral coherence according to previously described methods (41). The equation for spectral coherence is as follows:

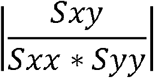

Where

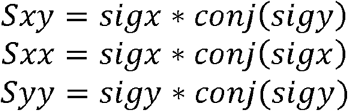

Spectral coherence was determined for the 10s window before and 10s after the start of a single drinking bout at least 5 seconds in duration (1 bout per condition per mouse). For statistical comparisons, the mean coherence in the 1 to 1.25 Hz frequency band was computed for each animal, and two-way ANOVA was used to compare coherence pre- and post-CA at each time window.

## Results

### BNST and CeA neurons exhibit acute calcium increases time-locked to drinking and eating behavior

We first assessed pan-neuronal activity during drinking behavior in C57BL/6J mice. Mice were injected in contralateral BNST and CeA with AAV5-hsyn-GcAMP6s, with optical fibers implanted above each injection site (Figure 1A). Optical fibers were located across predominantly dorsal BNST and throughout the CeA (Figure S1). Calcium activity in both regions transiently increased during bouts of alcohol drinking (Figure 2F,H). We also observed that the magnitude of this increase was attenuated with repeated alcohol drinking bouts, illustrated by the representative traces in Figure 2A and averaged traces in Figure 2D,E. In both regions, the mean calcium activity Z score during the drinking bout was lower in the last compared to first drinking bout in a session (2-5 bouts/mouse/session; Figure 2G,I).

**Figure 1.**
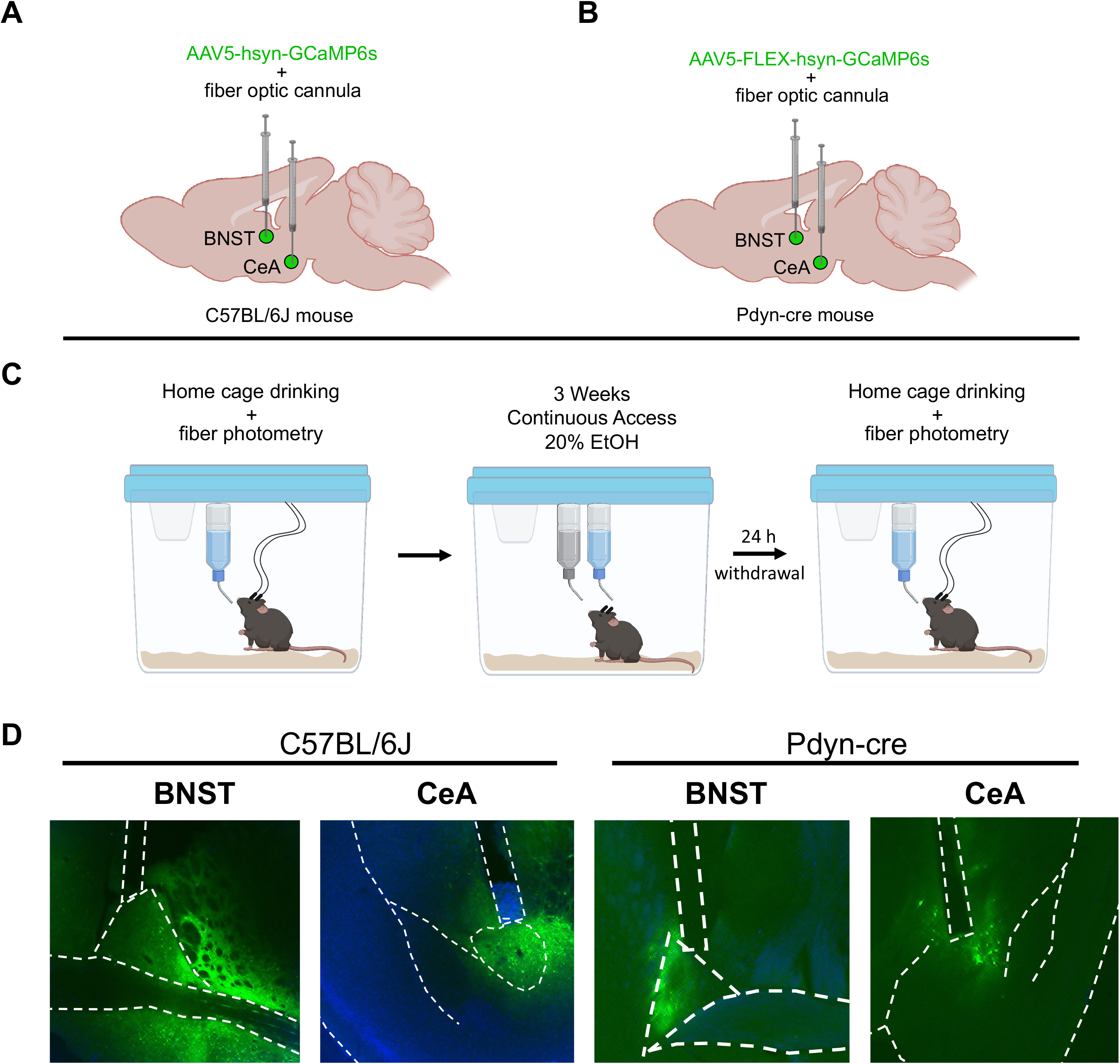
Overview of experimental design. (A) Wildtype (C57BL/6J) mice were stereotaxically injected with AAV5-hsyn-GCaMP6s to drive pan-neuronal expression of the calcium indicator GCaMP6s in the central amygdala (CeA) and contralateral bed nucleus of the stria terminalis (BNST) and implanted with optical fibers above each injection site. (B) Mice expressing cre under control of the preprodynorphin promoter (pdyn-cre) were injected with AAV5-hsyn-FLEX-GCaMP6s to target GCaMP expression to dynorphin-expressing neurons in the CeA and contralateral BNST with corresponding optical fiber placement. (C) Fiber photometry recordings of GCaMP in CeA and BNST were made during consumption of alcohol and other tastants in the home cage over 3-4 separate recording sessions. Mice were then provided with three weeks of continuous access to 20% w/v alcohol in the home cage. 24 hours following the end of continuous access, fiber photometry recordings were repeated to determine if chronic alcohol access affected calcium dynamics. (D) Representative images of virus and fiber placements in CeA and BNST in C57BL/6J and pdyn-cre mice.

**Figure 2.**
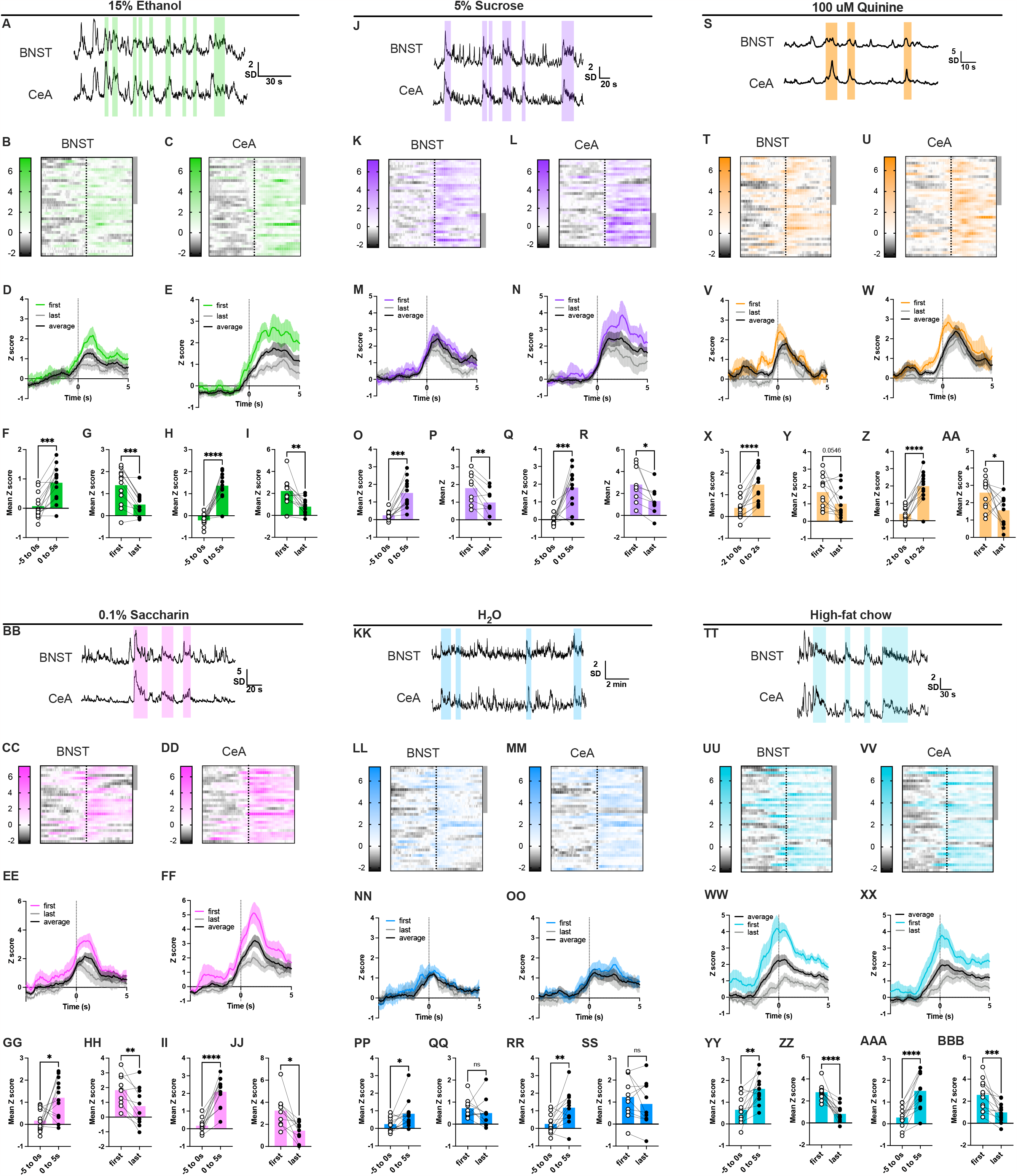
BNST and CeA neurons exhibit acute calcium increases time-locked to drinking and eating behavior. Data are shown for 15% ethanol (A-I), 5% sucrose (J-R), 100 μM quinine (S-AA), 0.1% saccharin (BB-JJ), water (KK-SS), and high-fat chow (TT-BBB). Representative traces from BNST and CeA demonstrate acute increases in activity during bouts of consumption of alcohol (A), sucrose (J), quinine (S), saccharin (BB), water (KK) and high-fat chow (TT) (bouts indicated by shading). Heatmaps show all 5s consumption bouts recorded in BNST and CeA for alcohol (B,C), sucrose (K,L), quinine (T,U), saccharin (CC,DD), water (LL, M), and high-fat chow (UU,VV) from n=13 mice, with the onset of each 5s bout aligned to the dotted line. Data from male mice (n=6) are indicated by grey shading on the right side of the heatmaps. Peri-event plots show the mean±SEM GCaMP6s activity during the first, last, and average consumption bouts in BNST and CeA for alcohol (D,E), sucrose (M,N), quinine (V,W), saccharin (EE,FF), water (NN,OO), and high-fat chow (WW,XX). Time zero represents the start of consumption bouts. For both regions, the mean Z score significantly increased during 5-second consumption bouts compared to the 5 seconds prior to bout onset for alcohol (F,H), sucrose (O,Q), quinine (X,Z), saccharin (GG,II), water (PP,RR), and high-fat chow (YY,AAA). (paired t-test, *p<0.05, **p<0.01, ***p<0.001, ****p<0.0001) (For quinine, drinking bouts were only 2 seconds in duration.) The mean Z score during consumption decreased from the first to last bout in BNST and CeA for alcohol (G,I), sucrose (P,R), quinine (Y,AA), saccharin (HH,JJ), and high-fat chow (ZZ,BBB) (paired t-test, *p<0.05, **p<0.01, ***p<0.001, ****p<0.0001). There was no change from the first to last bout for water for either region (QQ,SS).

To test whether these responses were specific to alcohol drinking, we tested other fluids of varying valence and caloric value. We found that sweet fluids, both caloric (5% sucrose, Figure 2O,Q) and non-caloric (0.1% saccharin, Figure 2GG,II), increased calcium activity, as did the aversive tastant quinine (100 μM, Figure 2X,Z), and water (Figure 2PP,RR). For all fluids except water, the mean Z score during drinking behavior decreased with repeated drinking bouts of the same fluid (Figure 2J,S,BB,KK), suggesting response amplitude is higher when fluids are novel. Finally, to test whether the calcium increase was specific to fluid consumption or to consummatory behavior more generally, we provided mice with access to a pellet of high-fat chow. Similar to fluid consumption, high-fat chow consumption was associated with an acute increase in calcium activity that was attenuated with repeated bouts (Figure 2YY-BBB). For all fluids, no sex differences were detected between males and females in two-way ANOVAs comparing the pre-vs. post-bout onset Z score; thus, data from both sexes were combined in Figure 2. Data from all statistical comparisons are included in Supplemental Tables.

We performed a subset of recordings without water deprivation to assess how the degree of thirst influences calcium activity during consumption of either alcohol or water (Figure S2). These recordings were made in the home cage but were one hour in duration to provide a longer opportunity to drink. There was no difference in the mean Z score in BNST or CeA during bouts of water or alcohol consumption compared to values obtained from the same mice following water deprivation (Figure S2F,H,N,P), suggesting that thirst is not the primary driver of the observed calcium responses. However, in the BNST the increase in Z score during water consumption without deprivation was not significant (Figure S2M), suggesting that in this region, thirst drives part of the response in water-deprived mice. Several mice also consumed standard mouse chow during these recordings; thus, we compared the signal obtained during standard chow consumption to that during high-fat chow consumption. Signal in CeA was not different between the two chow types (Figure S2X). However, the BNST showed a significantly reduced mean signal amplitude during chow versus high-fat food consumption (Figure S2V), and the change in Z score during standard chow consumption was not significant (Figure S2U), possibly reflecting differences in the novelty and reward value of the two food types.

### BNST and CeA dynorphin neurons exhibit acute calcium increases time-locked to drinking and eating behavior

We next assessed acute changes in calcium dynamics during drinking behavior in the subpopulation of dynorphin neurons in BNST and CeA. Male and female pdyn-cre mice were injected unilaterally in BNST and CeA with AAV5-hsyn-flex-GCaMP6s and optical fibers implanted in each region (Figure 1B). As in the previous cohort, fibers were localized to predominantly dorsal BNST and throughout the CeA (Figure S1). Similar to the response of the overall population in each region, calcium activity in dynorphin neurons was increased during bouts of drinking of all fluids tested, as well as during high-fat diet consumption, in CeA (Figure 3). In BNST, calcium activity was increased during the intake of high-fat diet and all fluids tested except for water. In contrast to pan-neuronal responses, in the dynorphin population we did not detect a significant change in the response from the first to last bout in a single drinking session for any fluid (Figure 3). We also detected significant sex differences for CeA responses to saccharin and quinine, where females had a greater mean response than males (see Supplemental Tables).

**Figure 3.**
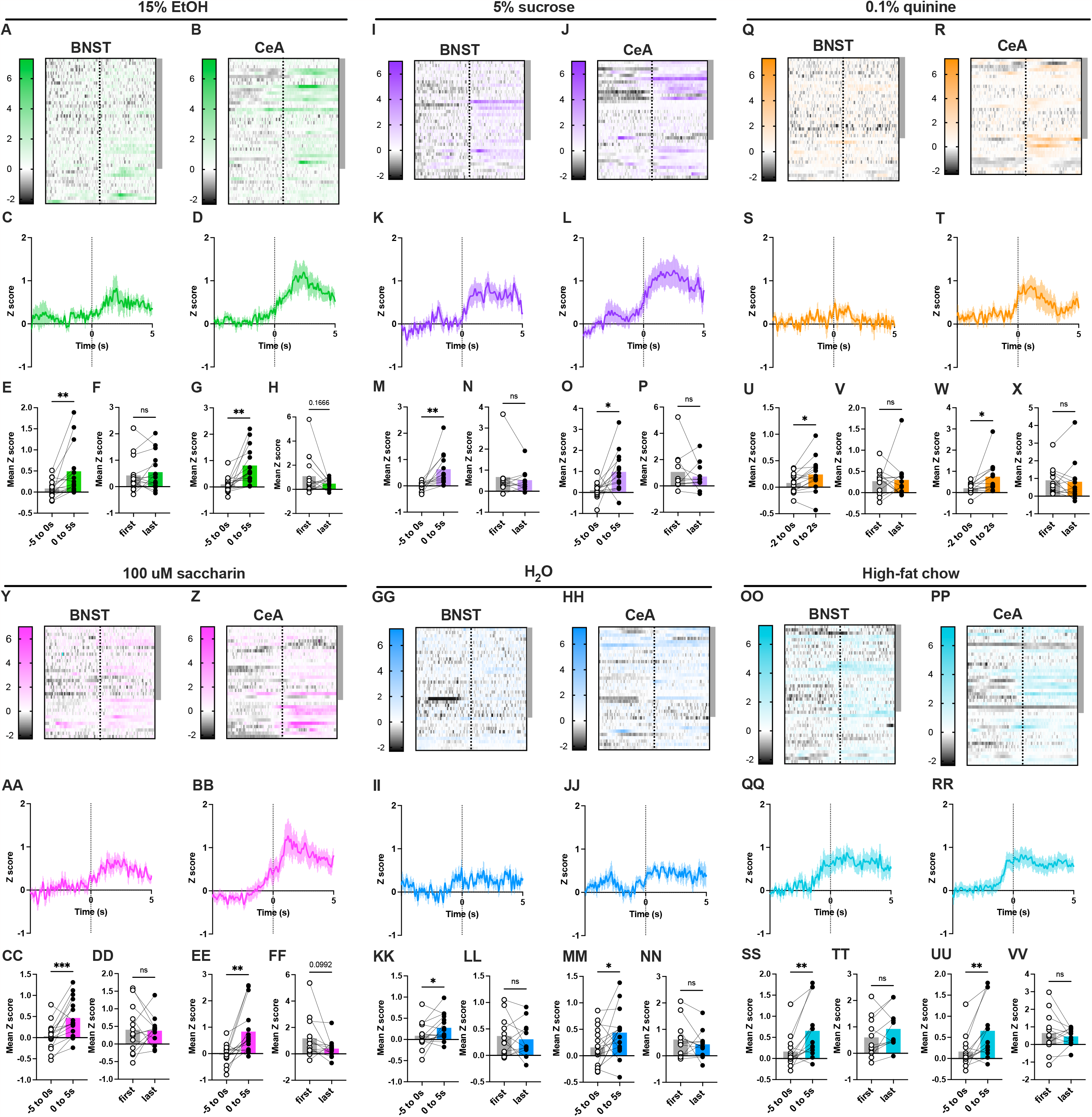
BNST and CeA dynorphin neurons exhibit acute calcium increases time-locked to drinking and eating behavior. Data are shown for 15% ethanol (A-H), 5% sucrose (I-P), 100 μM quinine (Q-X), 0.1% saccharin (Y-FF), water (GG-NN), and high-fat chow (OO-VV). Heatmaps show all 5s consumption bouts recorded in BNST and CeA for alcohol (A,B), sucrose (I,J), quinine (Q,R), saccharin (Y,Z), water (GG,HH), and high-fat chow (OO,PP) from n=16 mice, with the onset of each 5s bout aligned to the dotted line. Data from male mice (n=11) are indicated by grey shading on the right side of the heatmaps. Peri-event plots show the mean±SEM GCaMP6s activity during the first, last, and average consumption bouts in BNST and CeA for alcohol (C,D), sucrose (K,L), quinine (S,T), saccharin (AA,BB), water (II,KK), and high-fat chow (QQ,RR). Time zero represents the start of consumption bouts. The mean Z score significantly increased in BNST and CeA during 5-second consumption bouts compared to the 5 seconds prior to bout onset for alcohol (E,G), sucrose (M,O), quinine (U,W), saccharin (CC,EE), water (KK,MM), and high-fat chow (SS,UU) (paired t-test, *p<0.05, **p<0.01, ***p<0.001, ****p<0.0001). (For quinine, drinking bouts were 2 seconds in duration.) There was no change in mean Z score in the first compared to last bout of consumption for any tastant (F,H,N,P,V,X,CC,EE,LL,NN,TT,VV).

### BNST and CeA pan-neuronal activity during initial alcohol drinking bouts is reduced in females following 3 weeks of continuous access

We were interested in how chronic alcohol consumption would impact BNST and CeA dynamics during consummatory behaviors. To assess this, mice were provided with continuous access to 20% w/v alcohol for three weeks (Figure 1C). Female C57BL/6J mice drank significantly more alcohol and showed a greater ethanol preference than males of the same strain (Figure 4A,B). 24 hours following the removal of alcohol bottles, fiber photometry recordings were made to evaluate neuronal activity associated with drinking behavior during short-term alcohol withdrawal. Because of the attenuation of the signal with repeated bouts in the pan-neuronal population, average signal amplitude can be skewed based on the total number of bouts; thus, we compared first bouts between conditions. Twenty-four hours after the end of continuous access, in females we detected a significantly reduced change in the mean Z score from the pre-bout baseline in BNST, and a significantly reduced peak Z score in CeA, which were partially restored 3 weeks post-CA (Figure 4K,R). We also compared the first to last drinking bout for the subset of animals with multiple bouts meeting the criteria of ≥ 5s duration and ≥ 10s apart, with male and female subjects combined due to a limited n. Surprisingly, we found no difference in signal amplitude from first to last bout 24 h following CA, suggesting a loss of the novelty aspect of the response (Figure 4E,M). At 3 weeks post-CA, there was a restoration of the novelty response in CeA (Figure 4O); however, in BNST the mean Z during drinking bouts increased, rather than decreased, with successive drinking bouts (Figure 4G), demonstrating prolonged changes in how the BNST responds to alcohol following exposure to chronic drinking.

**Figure 4.**
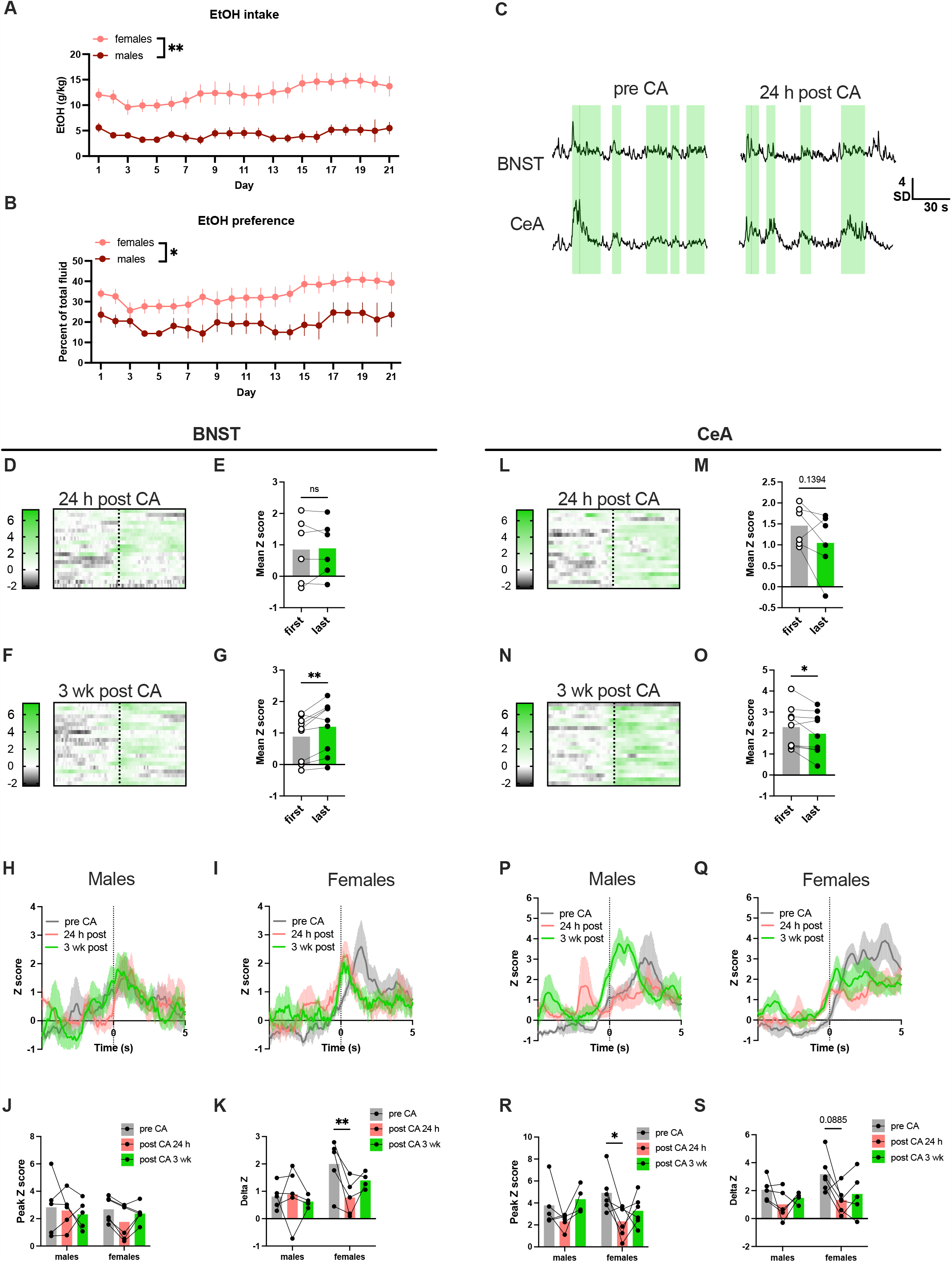
BNST and CeA pan-neuronal activity during initial alcohol drinking bouts is reduced in females following 3 weeks of continuous access (CA). Alcohol intake (A) and preference ratio (B) was significantly greater in female than male mice. (C) Representative traces from BNST (top) and CeA (bottom) showing acute increases in neuronal activity during bouts of alcohol consumption (indicated by green shading) 24 hours following the end of CA. (D,F) Heatmaps showing alcohol drinking bouts 24 hours (D) and 3 weeks (F) following the end of CA in BNST. (E) 24 hours post CA, there was no change in the mean Z score from the first to last bout of alcohol drinking in BNST in the subset of mice that drank multiple bouts. (G) 3 weeks post CA, there was an increase in the mean Z score from the first to last bout of alcohol drinking in BNST (paired t-test, **p<0.01). (H-I) Peri-event plots showing the mean±SEM GCaMP6s activity for average consumption bouts in male (H) and female (I) mice in BNST pre-CA (green), 24 hours post CA (red), and 3 weeks post CA (grey). Time zero represents the start of consumption bouts. (J) Following CA, there was no change in the peak Z score in BNST in either sex. (K) 24 hours post CA in females, the mean Z score increase from baseline during alcohol drinking bouts was smaller compared to pre-CA. (L,N) Heatmaps showing alcohol drinking bouts 24 hours (L) and 3 weeks (N) following the end of CA in CeA. (M) 24 hours post CA, there was no change in the mean Z score from the first to last bout of alcohol drinking in CeA in the subset of mice that drank multiple bouts. (O) 3 weeks post CA, there was a decrease in the mean Z score from the first to last bout of alcohol drinking in CeA (paired t-test, *p<0.05). (P,Q) Peri-event plots showing mean+-SEM GCaMP6s activity for average consumption bouts in male (P) and female (Q) mice in CeA pre-CA (green), 24 hours post CA (red), and 3 weeks post CA (grey). (R) 24 hours post CA, there was a main effect of CA (p<0.05) on the peak Z score, and in females, there was a significantly smaller peak Z score in CeA during alcohol drinking (mixed model ANOVA with Tukey post-hoc test, *p<0.05, **p<0.01). (S) 24 hours post CA, there was a main effect of CA and a trend for a reduction in the Z score change from baseline during acute alcohol drinking bouts in females in CeA.

At 24 h withdrawal, we also recorded BNST and CeA calcium activity during bouts of sucrose, water, and high-fat diet consumption to determine if the observed changes were specific to alcohol (Figure S3). Interestingly, during initial consumption bouts the calcium activity in BNST and CeA was lower during sucrose and water drinking, but not during high-fat diet consumption, as compared to pre-CA values (Figure S3G-J,Q-T). For most measures, these effects were specific to males.

In contrast to C57BL/6J mice, no sex differences in drinking behavior were detected in pdyn-cre mice (Figure S4A,B), who exhibited generally low levels of alcohol consumption for this paradigm. Because dynorphin neurons did not display attenuating responses across a session, we compared the mean Z scores during consumption bouts before and after CA (Figure S4). Dynorphin neurons in both regions remained responsive to the same stimuli in both BNST (Figure S4G,O,W,EE) and CeA (Figure S4I,Q,Y,GG), and we detected no changes relative to pre-CA (Figure S4H,J,P,R,X,Z,FF,HH), which likely reflects the low levels of alcohol consumption in these mice.

### BNST-CeA coherence is increased during alcohol drinking in males

Oscillatory synchronization is a key mechanism through which neural populations transmit information between regions (41), and previous studies have demonstrated changes in neurotransmission between CeA and BNST following chronic alcohol (37). To determine if BNST-CeA oscillatory coupling was altered during acute drinking bouts or following chronic drinking, we assessed the spectral coherence of the GCaMP signal simultaneously acquired from BNST and CeA during alcohol and sucrose consumption both before and after CA. We employed a previously established method (40) to calculate the spectral coherence, which computes the correlation between the two signals across a range of oscillatory frequencies. Figure 5 shows the BNST-CeA spectral coherence results for males and females in the 10s prior to and 10s following onset of alcohol (Figure 5A-D) and sucrose (Figure 5F-I) consumption, both before and after continuous access. Alcohol drinking-induced coherence changes were most prominent between 1-1.25 Hz, so we quantified mean coherence in this frequency range for statistical comparisons. Both before and after CA, males but not females showed increased BNST-CeA coherence during drinking bouts, and there was a main effect of CA to reduce overall coherence (Figure 5E). No changes in mean coherence between 1-1.25 Hz were observed for sucrose drinking either before or after CA (Figure 5J).

**Figure 5.**
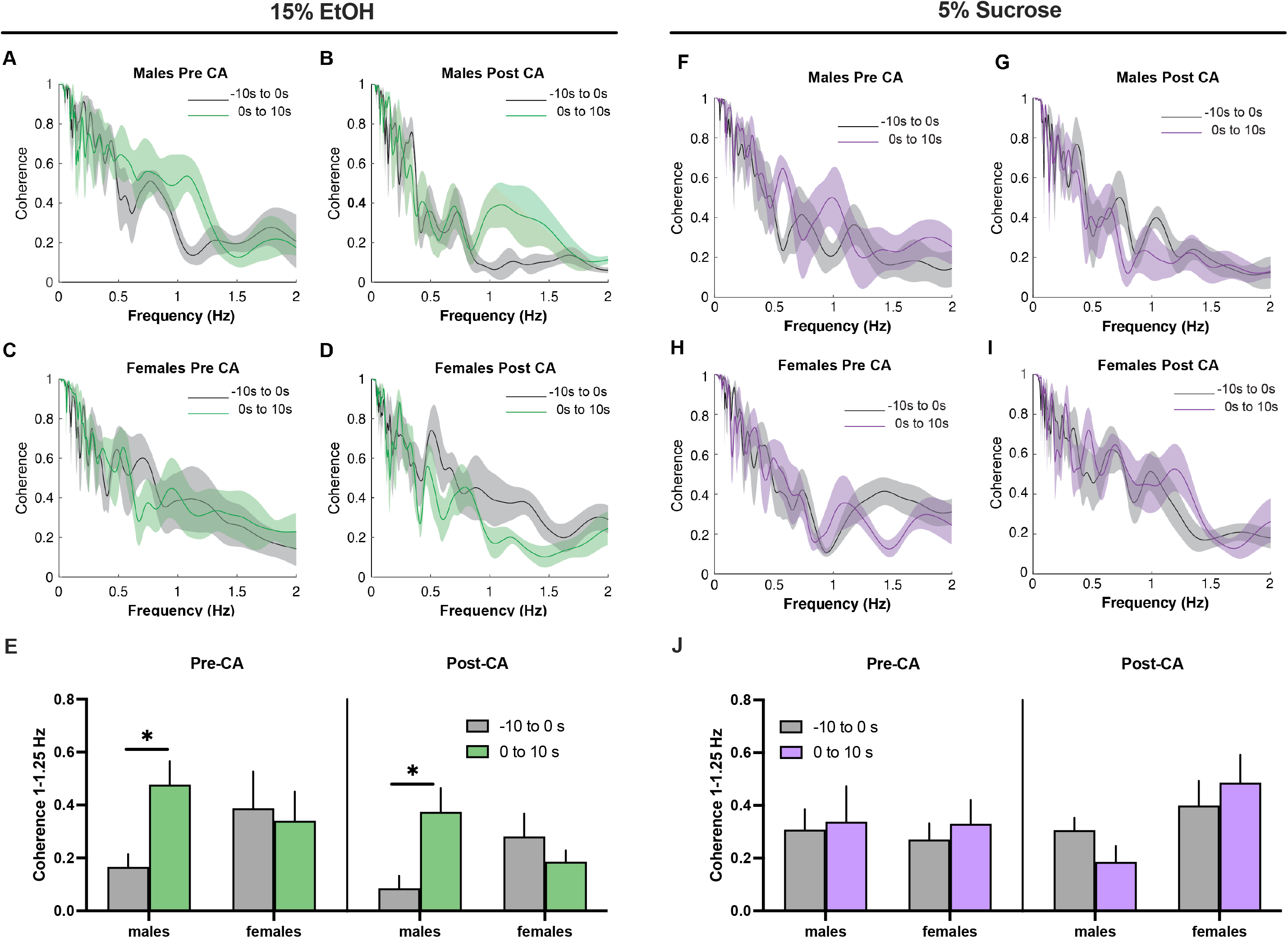
BNST-CeA coherence is increased during alcohol drinking in males. (A-D) Coherence spectra for 0-2 Hz for males pre-(A) and post-(B) continuous access (CA), and females pre-(C) and post-CA (D) for the 10-second period prior to alcohol drinking bouts (gray) and 10 seconds following bout onset (green). Alcohol drinking-induced coherence changes were greatest between 1 and 1.25 Hz. (E) BNST-CeA coherence between 1 and 1.25 Hz was increased in the 10 s post bout onset compared to the 10 s prior to bout onset for alcohol drinking in males both before and after CA. There was a main effect of CA to reduce coherence (3-way ANOVA with Sidak’s post hoc test, p<0.05). (F-I) Coherence spectra for 0-2 Hz for males pre-(F) and post-(G) continuous access (CA), and females pre-(H) and post-CA (I) for the 10-second period prior to sucrose drinking bouts (gray) and 10 seconds following bout onset (purple). (J) BNST-CeA coherence between 1 and 1.25 Hz was unchanged during sucrose drinking.

## Discussion

In this study, we used fiber photometry to examine dynamic activation and coherence of the BNST and CeA during home-cage drinking behavior before and after a continuous access drinking paradigm. Our findings revealed an increase in pan-neuronal and dynorphinergic BNST and CeA calcium activity during drinking behavior that was not specific to alcohol, as similar calcium changes were observed during the consumption of fluids of varying valence and caloric value, as well as high-fat food. Except for water, pan-neuronal responses were diminished, but not eliminated, with repeated consumption of the same tastant. This finding suggests that novelty influences neuronal engagement in BNST and CeA, and drinking-associated neuronal responses comprise both a transient novelty-sensitive component and a sustained component that may reflect reward. While dynorphin neurons showed similar increases in activity during consumption of nearly all fluids and food, they did not consistently display the same attenuating property with repeated bouts, indicating that dynorphin neurons contribute to the sustained but not novelty-driven component. Short-term withdrawal from continuous access eliminated the signal attenuation with repeated drinking bouts and reduced neuronal activation during the first drinking bout in females, suggesting that prior alcohol exposure reduces novelty associated with alcohol consumption. This change in signal could also reflect altered pharmacological properties of alcohol, such as tolerance, but our data do not allow us to disambiguate these factors. We also found that BNST-CeA coherence was increased during alcohol drinking bouts in males and reduced following continuous access, suggesting that both acute and chronic alcohol exposure change how these brain regions interact.

While the pan-neuronal approach yields insight into how brain regions are behaving on a larger scale, the BNST and CeA each comprise multiple subdivisions with diverse cell phenotypes and functions, and we are likely recording signal from multiple cell types (42). Indeed, while dynorphin neurons were activated during the same consummatory behaviors as BNST and CeA overall, their activity could not account for novelty sensitivity, and the cell types underlying this characteristic remain unknown. Identifying these populations will be an important area for future studies, as novelty reactivity of the BNST and amygdala in humans correlates with measures of anxiety (43,44). Signal attenuation with repeated intake of the same tastant may reflect reduced salience of the cue as it becomes less novel, or it may correspond to a reduction in arousal because the cue is established not to pose a threat, as novel fluids encountered in natural environments represent potential harms. One possible cell type mediating novelty sensitivity is PKCδ neurons, which are active in BNST during approach behavior in anxiogenic contexts (45,46) and play a role in fear behavior in CeA (47). Another possible mediator is CRF-expressing neurons, which are activated by aversive odors in BNST (13).

The effects of CA were specific to females, potentially due to their higher alcohol intake, with males showing similar directional changes in CeA that did not reach significance. Interestingly, males showed decreased responses to sucrose and water at 24 hours of withdrawal, which may represent a generalized reduction in salience or reward responsiveness, as withdrawal from even short-term alcohol exposure is associated with reduced responding to natural rewards (48). Responses to rewarding food, which may engage different pathways than fluids, were similar pre- and post-CA. Of note, previous studies have shown that different neuronal subpopulations are activated during the consumption of appetitive vs aversive fluids. For example, dopamine release in dorsolateral BNST increases during sucrose consumption, but decreases during quinine consumption, whereas norepinephrine release varies in the opposite direction (26,49). Thus, oppositely valenced stimuli may have opposing effects in different cell types that offset each other when recording bulk signal. In a similar vein, the suppression of BNST responses to sucrose and water 24 hours post CA in males are not necessarily absent from females but could be masked by an opposing change in a second population; for example, rewarding VTA projections might show reduced activity, but another population, such as CRF, might show increased engagement in females due to their higher alcohol intake.

Simultaneous activation of dynorphin neurons in BNST and CeA during fluid and food consumption is consistent with observations that neuropeptide-expressing cells in these regions generally play similar functional roles (3). These cells may be involved in signaling taste or orosensory reward, as suggested by a study in prodynorphin knockout mice (50). In this regard, there is extensive evidence for CeA involvement in taste processing, including in taste novelty (51), affective taste responding, and experience-dependent taste plasticity (52,53). In humans, the CeA controls a central gain mechanism influencing taste perception (54). As a stress-sensitive neuropeptide, dynorphin could be important for altering taste sensitivity in stressful contexts. However, a previous study showed that deletion of dynorphin or KOR from the CeA reduces ethanol intake without affecting the intake of sweet or bitter fluids (55), suggesting that taste modulation is not the primary mechanism by which CeA dynorphin influences alcohol drinking. An important caveat to our observations is that dynorphin-expressing cells can co-express additional neurotransmitters, including GABA, CRF, and somatostatin, and we do not know which transmitters are released during drinking behavior. Although a role for dynorphin and KOR in alcohol drinking is well established, a measurement of *in vivo* dynorphin release, e.g., using a dynorphin sensor, would be needed to establish dynorphin release in this drinking paradigm. Another possibility is that these neurons release GABA during drinking behavior under basal conditions, and during stress, dynorphin is co-released to activate presynaptic KORs, suppressing GABAergic transmission (56). This could serve as a potential mechanism for stress-induced dampening of consummatory behaviors or consumption-associated reward.

Importantly, we did not study intoxication in the present study. During fiber photometry recordings, alcohol bottles were removed after several drinking bouts and recording sessions terminated shortly thereafter in order to study the acute response associated with consumption rather than intoxication. Electrophysiology experiments indicate that acute application of ethanol to slices increases inhibitory neurotransmission in CeA and BNST (20,57), suggesting that these regions should become inhibited at high blood alcohol concentrations. The effects of intoxication on BNST and CeA activity in vivo remains an important question for further study. Additionally, we used a continuous access protocol that generates stable but non-escalating patterns of drinking and does not produce alcohol dependence. Higher levels of alcohol consumption could generate differential changes due to the engagement of dependence-related circuitry. Given that pdyn-cre mice in this study voluntarily consumed low quantities of alcohol, and we were unable to detect any changes in dynorphin neuron activity post CA, vaporized ethanol may be a better paradigm for future studies.

This study is the first to assess BNST-CeA coherence using fiber photometry, an emerging application for this type of data (40,58). Compared to electrophysiological methods, the temporal resolution of GCaMP is lower and permits measurement of oscillatory activity at relatively low frequencies, which have not yet been correlated with specific neurotransmitter activities or behavioral states. However, GCaMP activity is correlated with local field potentials (59), spiking events in pEEGs (60), and fMRI signal fluctuations (40), albeit with different temporal dynamics. Our observation of increased coherence during alcohol drinking bouts in males, and reduced coherence following CA, are consistent with reports showing that alcohol can alter neurotransmission between these regions. CeA projections to the BNST show changes in inhibitory control following alcohol exposure, leading to increased CRF output to the BNST (37), and these projections drive alcohol withdrawal drinking (29). Intriguingly, BNST-CeA functional connectivity is correlated with measures of anxiety in humans (61,62), including in individuals with AUD during early abstinence (63). CeA to BNST circuits have also been implicated in anxiety-like behaviors in rodents (64). Thus, the coherence changes we detected may serve as a biomarker for behavioral pathology associated with alcohol exposure. These findings further suggest that alcohol acutely activates a common upstream driver of activity in BNST and CeA that is shaped by inhibitory contacts within and between the structures. Future studies are needed to determine how we can experimentally manipulate BNST, CeA, or shared inputs to produce behaviorally relevant changes GCaMP oscillatory synchrony.

## Acknowledgments

We acknowledge Dr. Meghan Flanigan, Dr. Sofia Neira, and Dr. J.R. Haun for technical assistance, and Dr. Brianna George and Sarah Chong for editing the manuscript.

## Author contributions

A.V.R. designed and performed experiments, analyzed data, and wrote the manuscript. S.N.M., T.R.S., and S.I.L. performed experiments. O.J.H., T.H.C., and Y.I.S. analyzed data. T.L.K. designed experiments and wrote the manuscript.

## Funding

This work was supported by the National Institutes of Health (NIH) National Institute of Alcohol Abuse and Alcoholism (NIAAA) grant U01AA020911and P60AA011605 to T.L.K., and T32AA007573 to A.V.R.

## Competing Interests

The authors have nothing to disclose.

## Notes

### Competing Interest Statement

The authors have declared no competing interest.

