## Supplemental Figures for "Acute and chronic alcohol modulation of extended amygdala calcium dynamics"

**Figure S1. Optical fiber placements in BNST and CeA.** Atlas location of optical fiber tips for all C57BL/6J and pdyn-cre mice in the indicated anterior-posterior plane relative to Bregma.

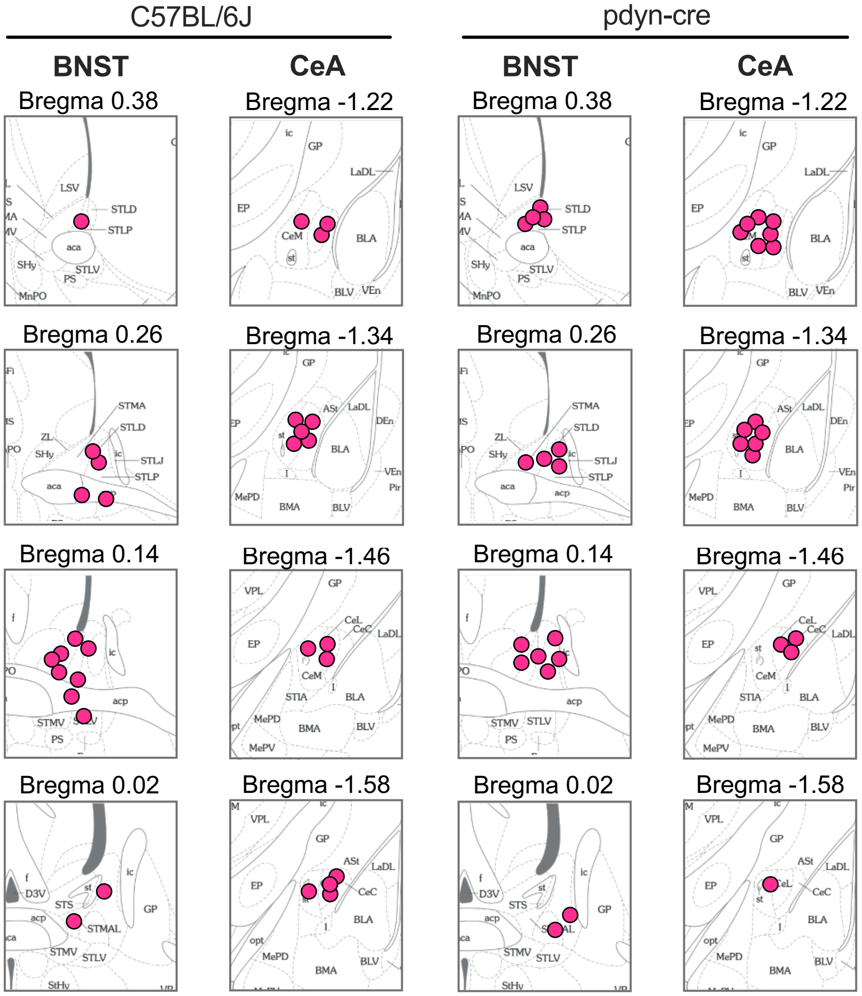

**Figure S2. BNST and CeA responses to alcohol are unchanged in the absence of water deprivation.** Data are shown for 15% ethanol (A-H), water (I-P), and food (Q-X). Heatmaps show all 5s consumption bouts recorded in BNST and CeA for alcohol (A,B), water (I,J), and standard chow (Q,R) from n=9 mice (5 male, 4 female) with the onset of each 5s bout aligned to the dotted line. Peri-event plots show the mean±SEM GCaMP6s activity in BNST and CeA during alcohol (C,D) and water (K,L) consumption under conditions of water deprivation (gray) or no water deprivation (green/blue); and during consumption of high-fat chow (HFD, blue) compared to standard chow (orange) (S,T). Time zero represents the start of consumption bouts. The mean Z score significantly increased during 5-second consumption bouts compared to the 5 seconds prior to bout onset in BNST and CeA for alcohol (E,G), and in CeA but not BNST for water (M,O) and standard chow (U,W) (paired t-test, *p<0.05, **p<0.01). There was no difference in the responses of BNST and CeA to alcohol (F,H) or water (N,P) during water deprivation compared to non-water-deprivation conditions. However, BNST (V) but not CeA (X) showed a reduced response to standard chow compared to regular chow (paired t-test, **p<0.01).

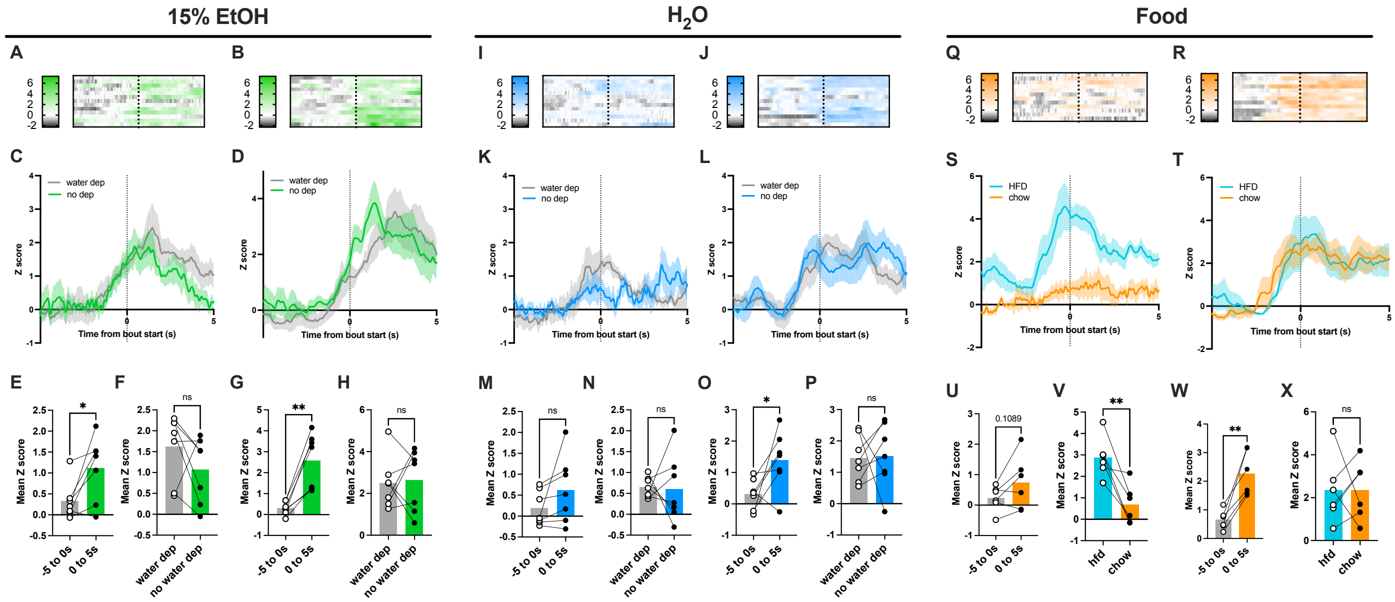

**Figure S3. Calcium activity during consumption of non-alcohol fluids is reduced 24 hours following continuous alcohol access in males.** Heatmaps show all consumption bouts for sucrose (A,B), water (K,L), and high-fat chow (U,V) 24 hours post continuous access (CA). (C-F) Peri-event plots show the mean±SEM GCaMP6s activity for average sucrose consumption bouts in male (C,D) and female (E,F) mice in BNST and CeA pre-CA (purple) and 24 hours post CA (red). (G) 24 h post CA, there was a main effect of CA to reduce the Z score change from baseline in BNST during sucrose drinking bouts. (H) 24 h post CA, there was a main effect of CA to reduce the Z score change from baseline in CeA during sucrose drinking bouts, with a trend for reduction in males. (I) 24 h post CA, there was a main effect of CA ($p<0.05$) on peak Z score in BNST during sucrose drinking, with a trend for reduction in males. (J) 24 h post CA, there was a main effect of CA and sex*CA interaction ($p<0.05$) on peak Z score, with a significantly smaller peak Z score in CeA during sucrose drinking in males but not females. (M-P) Peri-event plots show the mean±SEM GCaMP6s activity for average water consumption bouts in male (M,N) and female (O,P) mice in BNST and CeA pre-CA (blue) and 24 hours post CA (red). (Q) 24 h post CA, there was a main effect of CA to reduce the Z score change from baseline in BNST during water drinking bouts (R) 24 h post CA, there no effect of CA on the  Z score change from baseline in CeA during water drinking bouts. (S) 24 h post CA, there was no effect of CA on peak Z score in BNST during water drinking. (T) 24 h post CA, there was a main effect of CA and sex*CA interaction ($p<0.05$) on peak Z score, with a significantly smaller peak Z score in CeA during sucrose drinking in males but not females. (W-Z) Peri-event plots show the mean±SEM GCaMP6s activity for average food consumption bouts in male (W,X) and female (Y,Z) mice in BNST and CeA pre-CA (teal) and post-CA (red). There were no effects of CA on either the change in Z score from baseline (AA,BB) or peak Z score (CC,DD) in either region.

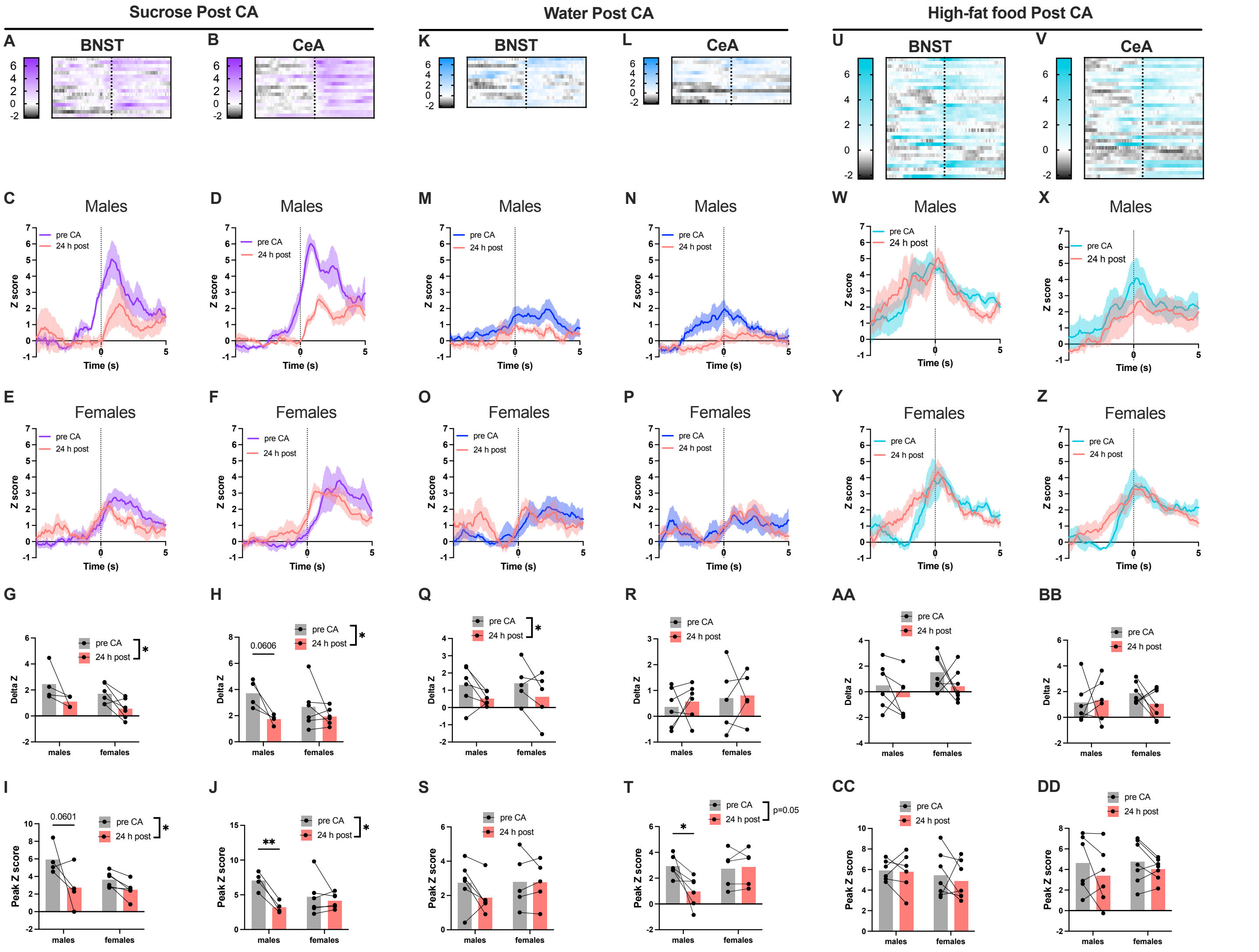

**Figure S4. Following continuous access, BNST and CeA dynorphin neurons exhibit no change in acute calcium increases time-locked to drinking and eating behavior.** Alcohol intake (A) and preference ratio (B) was not different between male and female mice. Fiber photometry data are shown for 15% ethanol (C-J), 5% sucrose (K-Q), water (S-Z), and high-fat chow (AA-HH). Heatmaps show all 5s consumption bouts recorded in BNST and CeA for alcohol (C,D), sucrose (K,L), water (S,T), and high-fat chow (AA,BB) from n=16 mice, with the onset of each 5s bout aligned to the dotted line. Peri-event plots show the mean±SEM GCaMP6s activity in BNST and CeA for consumption of alcohol (E,F), sucrose (M,N), water (U,V), and high-fat chow (CC,DD). Time zero represents the start of consumption bouts. The mean Z score significantly increased in BNST and CeA during 5-second consumption bouts compared to the 5 seconds prior to bout onset for alcohol (E,I), sucrose (O,Q), and high-fat chow (EE,GG), but only in CeA for water (W,Y) (paired t-test, $*p<0.05$, $**p<0.01$, $***p<0.001$). There was no change in the mean Z score during consumption pre- and post-CA for any tastant (H,J,P,R,X,Z,FF,HH).

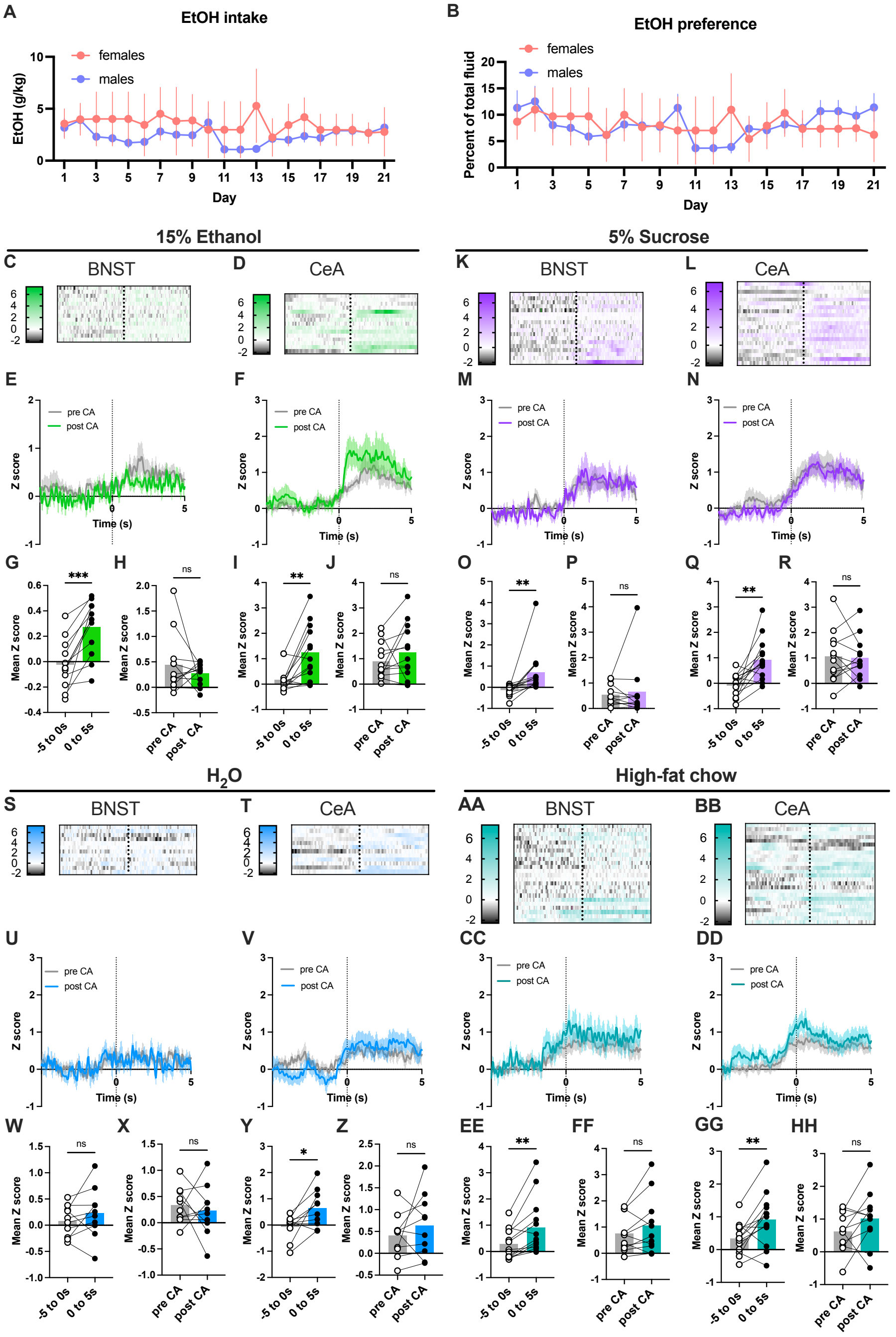
